# Quantifying Myelin Water Fraction in a Guinea Pig Model of Spontaneous Intrauterine Growth Restriction

**DOI:** 10.1101/2023.07.25.550586

**Authors:** Simran Sethi, Lanette J. Friesen-Waldner, Timothy R.H. Regnault, Charles A. McKenzie

## Abstract

Intrauterine growth restriction (IUGR) is an obstetrical outcome where a fetus has not achieved its genetic potential. A consequence of IUGR is a decrease in brain myelin content. Myelin water imaging (MWI) has previously assessed fetal myelin water fraction (MWF) and can potentially assess myelination changes associated with IUGR. Thus, this study aims to quantify and compare the MWF of non-IUGR and IUGR fetal guinea pigs (GPs) in late gestation. Our sample consisted of 22 pregnant Dunkin-Hartley GPs with 71 fetuses (34 male) [mean ± standard deviation: 60 ± 1.2 days gestation]. Eight SPGR volumes [flip angles (α): 2° – 16°], and two sets of 8 bSSFP volumes (α: 8° – 64°), at 0° and 180° phase increments were acquired at 3.0 T. MWF maps were generated for each fetal GP brain using multicomponent driven equilibrium single pulse observation of T_1_/T_2_ (mcDESPOT). Regions of interest (ROIs) were placed in the fetal corpus callosum (CC), fornix (FOR), and parasagittal white matter (PSW). Linear regression was performed between five fetal IUGR markers [body volume (BV), body-to-pregnancy volume ratio (BPrVR), brain-to-liver VR (BLVR), brain-to-placenta VR (BPlVR), and brain-to-BVR (BBVR)] and MWF for all regions (coefficient of determination, R^2^). A t-test with a linear mixed model compared the MWF of non-IUGR and IUGR fetal GPs for all three regions (α = 0.05). The MWF values are as follows: (mean ± standard deviation): 0.23 ± 0.02 (fetal CC), 0.19 ± 0.02 (fetal CC – IUGR), 0.31 ± 0.02 (fetal FOR), 0.27 ± 0.01 (fetal FOR – IUGR), 0.28 ± 0.02 (fetal PSW), and 0.24 ± 0.03 (fetal PSW – IUGR). Significant differences in MWF were found between the non-IUGR and IUGR fetuses in every region. In conclusion, the mean MWF of IUGR fetal GPs is significantly lower than non-IUGR fetal GPs.

## INTRODUCTION

Intrauterine growth restriction (IUGR) is an obstetrical complication where a fetus exhibits diminished growth below its genetic weight and size potential (1). According to the World Health Organization, a fetus with IUGR has an estimated weight below the 10^th^ percentile for their gestational age and a cut-off birth weight of 2500 grams at birth (2). IUGR is a leading cause of perinatal morbidity and mortality, second only to prematurity (3). The incidence of IUGR is estimated to be approximately 3-10% in the general obstetric population, although the incidence varies depending on the analyzed population, geographic location, and the standard growth curves used as a reference (4). Most cases of IUGR are due to an inadequate supply of oxygen and nutrients to the fetus due to placental insufficiency (5).

In humans, the neurological consequences of IUGR manifest as early as one year of age and persist throughout adolescence into adulthood; these neurological effects range from reduced cognitive skills and learning disabilities to neurological disorders such as autism spectrum disorder and cerebral palsy (6). The likelihood of the consequence depends on the insult’s length, severity, and timing. A significant factor in the pathogenesis of these disabilities is white matter injury characterized by multiple factors, including impaired myelination, as notable decreases in myelin content are seen between control newborn infants and those with IUGR (7).

Magnetic resonance imaging (MRI) is increasingly used for clinical and research purposes to assess fetal neurodevelopment (8). Myelin water imaging (MWI) is an MRI-based technique that images the aqueous components associated with the myelin sheath, specifically myelin water and intra-/extra-cellular water, to quantify myelin water fraction (MWF) (9). MWF is the ratio of the signal from myelin water to the signal from both myelin and intra-/extra-cellular water and is validated as a strong marker for myelin lipid (9). The feasibility of MWI has been successfully demonstrated in numerous animal models (10) and the fetal environment by quantifying MWF in a fetal guinea pig model (11). Preliminary results from the guinea pig work suggested that the MWF in the fetal corpus callosum and the fornix was lower in the fetuses with IUGR than in control fetuses (11). This current study builds on the pilot study by quantifying the difference in MWF in various brain regions. Guinea pigs are a suitable pre-clinical model of neurologic developmental programming for humans (12,13) and, like humans, can develop spontaneous IUGR (14,15). Thus, this study aims to determine whether MWF in various fetal brain regions differs in fetal guinea pigs with spontaneous IUGR compared to control guinea pigs. It was hypothesized that MWF would be lower in the fetuses with IUGR than those without IUGR.

## MATERIALS AND METHODS

The institution’s Animal Care and Ethics Committee reviewed, approved, and monitored the imaging protocol. The sample consisted of twenty-two pregnant, chow-fed female in-house bred Dunkin-Hartley guinea pigs (Charles River Laboratories) late in gestation (59-64 days gestation, term = ∼68 days) with a total of 73 fetuses. A spontaneous model of IUGR was used where IUGR occurred naturally without artificial interventions (14,15).

### Imaging

Before imaging, the guinea pigs were induced with 4.5% isoflurane with 2 litres/minute O_2_ and maintained via nose cone with a 1.5-2.5% isoflurane with 2 litres/minute O_2_. All imaging was performed with a 3.0 T MRI scanner (Discovery MR750, GE Healthcare, Waukesha, WI) with a 32-element human cardiac coil array (In Vivo Corp., Gainesville, FL). Anatomical T_2_-weighted ^1^H images of the entire uterus, including all fetal guinea pig brains, were acquired with a 3D spin-echo sequence: repetition time (TR)/echo time (TE) = 2002/218 ms, NEX = 2, field of view (FOV) = 10-12 cm, voxel size = 0.6 x 0.6 x 0.6 mm; the T_2_-weighted acquisition was approximately 10 minutes long.

Eight spoiled gradient echo (SPGR) volumes (TR/TE: 6.50/3.03 ms) for driven equilibrium single pulse observation of T_1_ (DESPOT_1_) were acquired at varying flip angles (α: 2° – 16°, increasing increments of 2°) in one acquisition (16). Two sets of eight balanced steady-state free precession (bSSFP) volumes (TR/TE: 6.7/3.4 ms) for DESPOT_2_ were acquired at various flip angles (α: 8° – 64°, increasing increments of 8°) and at 0° and 180° phase increments in a separate, single acquisition (16). The eight DESPOT_1_ and 16 DESPOT_2_ volumes were acquired in approximately 13 and 30 minutes, respectively. All DESPOT_1/2_ acquisitions had the following imaging parameters: NEX = 1, FOV = 10-12 cm, voxel size = 0.6 x 0.6 x 0.6 mm, acceleration factor: 2x [ASSET (DESPOT_1_) & ARC (DESPOT_2_)].

After imaging, the sows were monitored and kept on O_2_ until awake. The sows were transferred to be kept under a heating lamp and monitored until they were fully awake and mobile. The sows were then returned to their cages.

### MWF Map Reconstruction and MWF Quantification

Masks of the fetal guinea pig brains were manually generated from the T_2-_weighted images using FSLeyes (17). Using Quantitative Imaging Tools (18), multicomponent DESPOT_1/2_ (mcDESPOT) was used to reconstruct the MWF maps of each fetal guinea pig’s brain (19).

A trainee in medical imaging (6 years of experience) used 3D Slicer (v. 4.11.0-2019-12-02) (20–22) to place four to six regions of interest (ROIs) with a diameter of 0.5 mm and 1-2 pixels in the corpus callosum (CC), fornix (FOR) and parasagittal white matter (PSW) of each fetal guinea pig brain. The ROIs were placed on the T_2_-weighted images and then transferred to the MWF maps to quantity the regions’ MWF values. Images from Sethi *et al.* and Gareau *et al.* were used for reference to identify the CC with the FOR and PSW located below and above the CC, respectively (11,23). The ROI placements were done blinded and before IUGR determination.

### Animal Collection

The maternal sows were euthanized via CO_2_ inhalation 2-3 days after imaging (61-66 days gestation). All fetuses were removed from the sow, confirmed dead, and weighed immediately. Fetal brains, fetal livers, and placentae were removed and weighed immediately. The following IUGR weight markers and their IUGR cut-off values are commonly used in animal studies for IUGR determination: body weight (BW), body weight of each fetus to the average weight of the fetuses within the pregnancy ratio (BPrWR), brain-to-liver weight ratio (BLWR), brain-to-placenta weight ratio (BPlWR), and brain-to-body weight ratio (BBWR) (2,24–27). The IUGR cut-off value for each marker is as follows: BW ≤ 85 grams, BPrWR ≤ 0.9, BLWR ≥ 0.6, BPlWR ≥ 0.6, and BBWR ≥ 0.03.

### IUGR Determination

Although weight markers are commonly used in animal studies for IUGR determination, this study employed each weight marker’s corresponding volume marker instead for human translational purposes. Furthermore, the strong correlation between weight and volume demonstrated in previous work further supported the use of volume markers (11). Hence, fetal body, brain, liver, and placental volumes were quantified using the T_2_-weighted images. The corresponding five fetal volume markers were quantified for IUGR determination: body volume (BV), body volume of each fetus relative to the average volume of the fetuses within the pregnancy ratio (BPrVR), brain-to-liver volume ratio (BLVR), brain-to-placenta volume ratio (BPlVR), and brain-to-body volume ratio (BBVR).

Linear regression was performed between each weight marker and its respective volume marker to acquire the linear equation describing the correlation between each weight and its respective volume marker, resulting in five different linear equations. The IUGR cut-off value for each volume marker was then calculated using the above-mentioned IUGR cut-off value for its respective weight marker and linear equation. For instance, the BV cut-off value was calculated using the BW cut-off value, *i.e.,* 85 grams, and the linear equation describing the correlation between BW and BV. The calculated IUGR cut-off values for the volume markers are as follows: BV ≤ 37 cm^3^, BPrVR ≤ 0.9, BLVR ≥ 0.6, BPlVR ≥ 0.42, and BBVR ≥ 0.068. A fetus was determined to have IUGR if it met at least three of the five volume markers.

### Statistical Analysis

A simple linear regression analysis was performed between the following for all three regions separately: BV v. MWF, BPrVR v. MWF, BLVR v. MWF, BPlVR v. MWF, and BBVR v. MWF. F-tests were also conducted to determine if the slopes of the above-mentioned comparisons for all three regions were significantly different from zero (α = 0.05). Incorporating a linear mixed model, a one-way analysis of variance (ANOVA) was performed to compare the mean MWF of non-IUGR and IUGR fetuses for all three analyzed regions, with the sex of the fetus being a co-variate (α = 0.05). The linear mixed model was incorporated to consider litter effects as fetuses within a litter experience genetic and environmental similarities and, hence, are not independent (28).

The one-way ANOVA analysis incorporating the linear mixed model was performed using RStudio Team (2020). Rstudio: Integrated Development for R. Rstudio, PBC, Boston, MA URL http://www.rstudio.com/. The simple linear regression analysis and F-tests were performed using GraphPad Prism version 9.3.1 for Windows, GraphPad Software, San Diego, California, USA, www.graphpad.com.

## RESULTS

No adverse events occurred in the maternal guinea pigs during imaging. MWF maps were successfully generated for each fetal guinea pig’s brain. Two of the 73 fetal guinea pigs imaged were deceased at collection; data from those two fetuses were excluded. Of the 71 fetal guinea pigs analyzed, 19 were determined to have IUGR as determined by meeting at least three of the five criteria, comprising 27% of the sample (Tables 1 & 2).

**Table 1.**
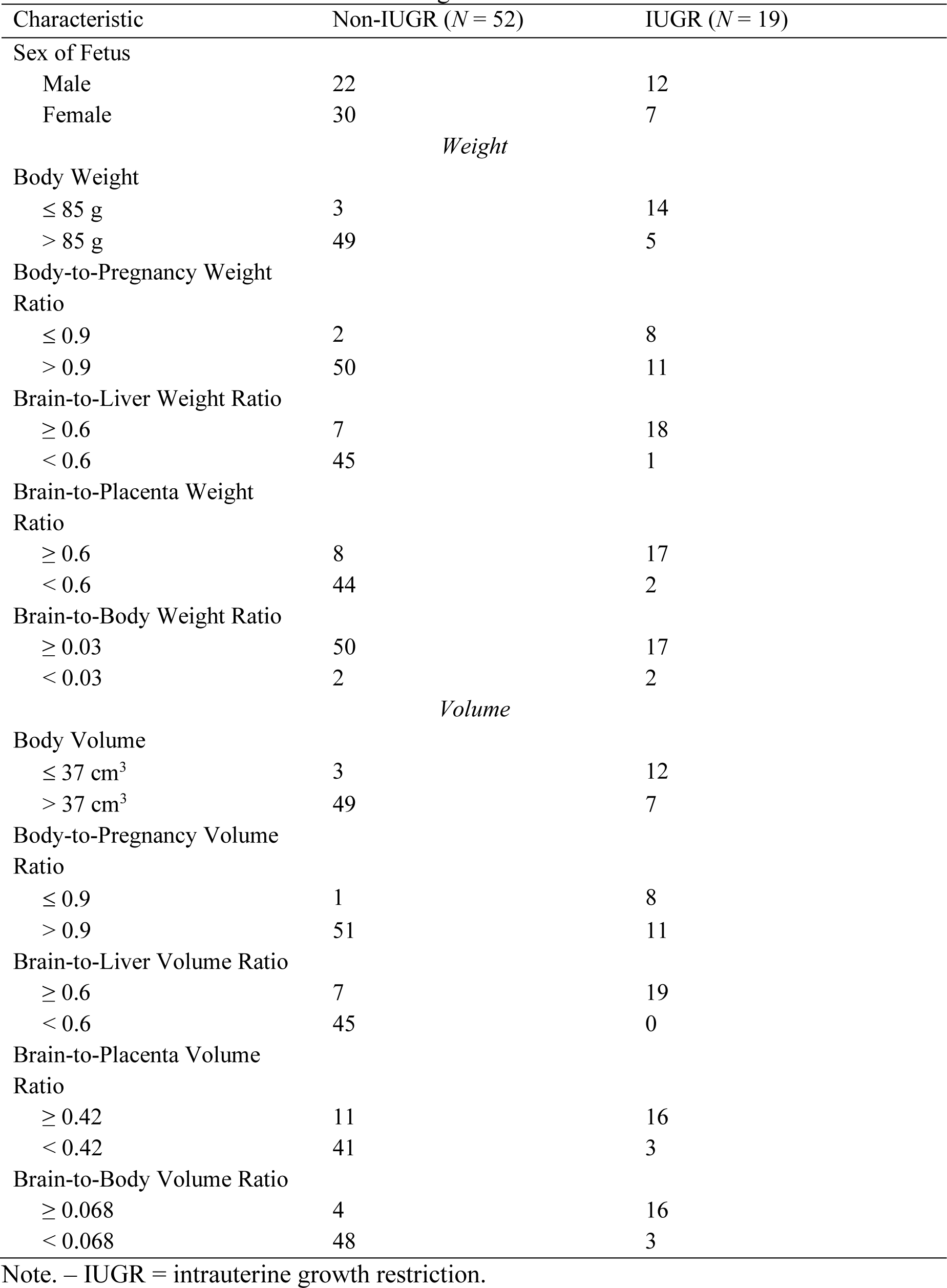
Characteristics of Fetal Guinea Pigs.

**Table 2.** Distribution of Fetal Guinea Pigs Within Maternal Guinea Pigs.

| Maternal Guinea Pig | Non-IUGR ($N = 52$) | IUGR ($N = 19$) |
| --- | --- | --- |
| 1 | 1 | 0 |
| 2 | 1 | 2 |
| 3 | 3 | 2 |
| 4 | 3 | 0 |
| 5 | 2 | 2 |
| 6 | 2 | 0 |
| 7 | 3 | 1 |
| 8 | 2 | 0 |
| 9 | 1 | 2 |
| 10 | 2 | 1 |
| 11 | 3 | 0 |
| 12 | 1 | 2 |
| 13 | 3 | 0 |
| 14 | 4 | 0 |
| 15 | 4 | 0 |
| 16 | 3 | 1 |
| 17 | 4 | 1 |
| 18 | 3 | 2 |
| 19 | 3 | 0 |
| 20 | 0 | 2 |
| 21 | 2 | 1 |
| 22 | 2 | 0 |

Note. – Maternal guinea pig #22 had two fetuses that were deceased before collection and are not reported in the table; IUGR = intrauterine growth restriction.

The linear regression results between each weight and its respective volume IUGR marker are shown in Table 3. The one-way ANOVA showed the mean MWF of the IUGR guinea pigs to be significantly lower than the mean MWF of non-IUGR guinea pigs for all three regions (*p* < 0.05) [Figures 1 & 2]. The simple regression results for all three regions between MWF and each IUGR volume marker are shown in the following figures: BV v. MWF (Figure 3), BPrVR v. MWF (Figure 4), BLVR v. MWF (Figure 5), BPlVR v. MWF (Figure 6), and BBVR v. MWF (Figure 7). The F-test results showed that the slope between MWF and each IUGR volume marker for every region analyzed significantly differed from zero (*p* < 0.05).

**Figure 1.**
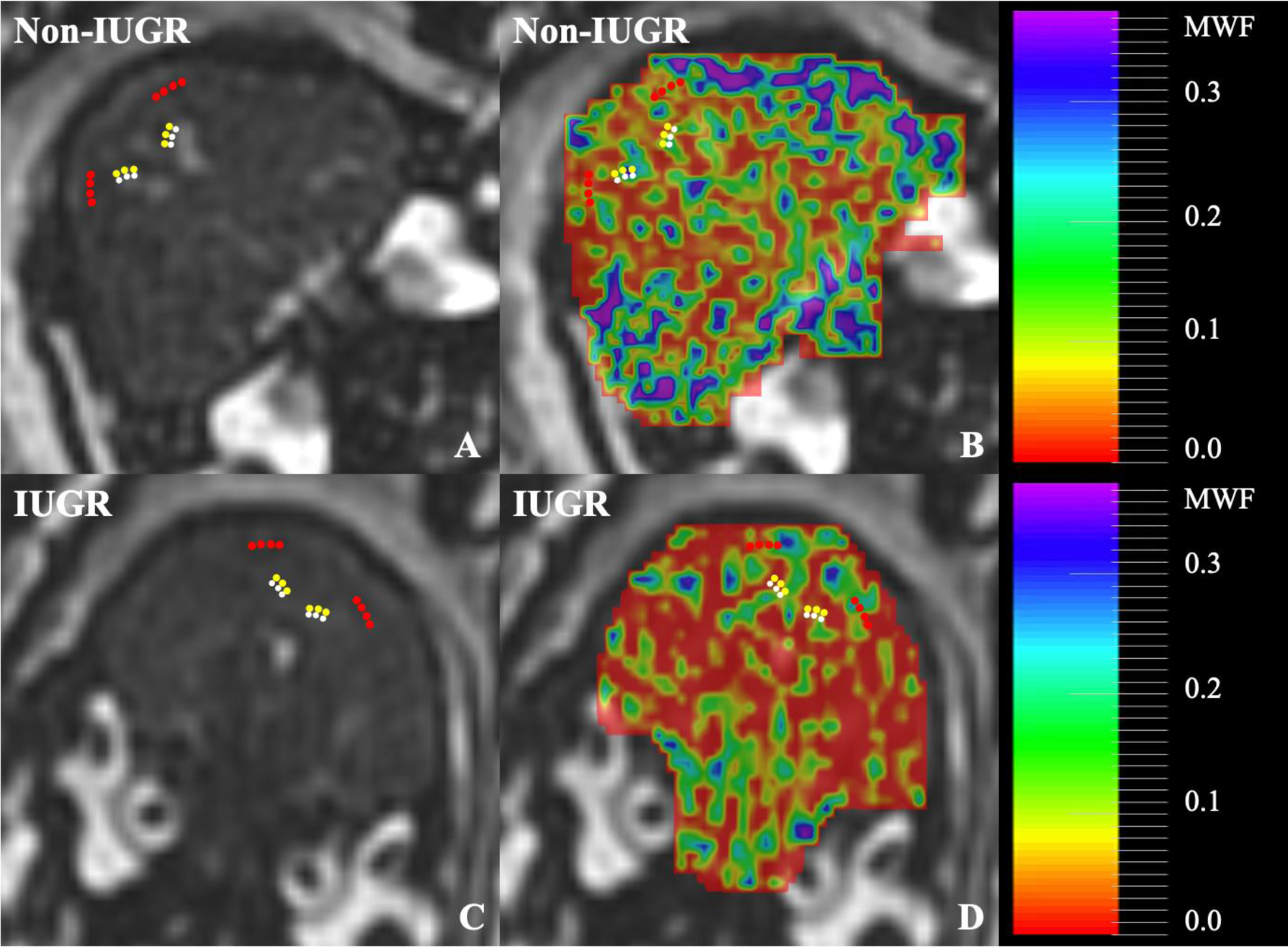
**(A)** Coronal T_2_-weighted image of a fetal guinea pig brain with no intrauterine growth restriction (IUGR) at 62 days gestation. **(B)** A 50% transparent myelin water fraction (MWF) map of the same guinea pig brain overlaid on the corresponding anatomical image shown in A. The fetus did not meet any of the IUGR criteria. **(C)** Coronal T_2_-weighted image of a fetal guinea pig brain with IUGR at 61 days gestation. **(D)** A 50% transparent MWF map of the same guinea pig brain overlaid on the corresponding anatomical image in C. The fetus met four of the five IUGR criteria. A colour scale for the MWF maps is shown for reference. In all four figures, regions of interest are placed in the corpus callosum (yellow), fornix (white), and parasagittal white matter (red).

**Figure 2.**
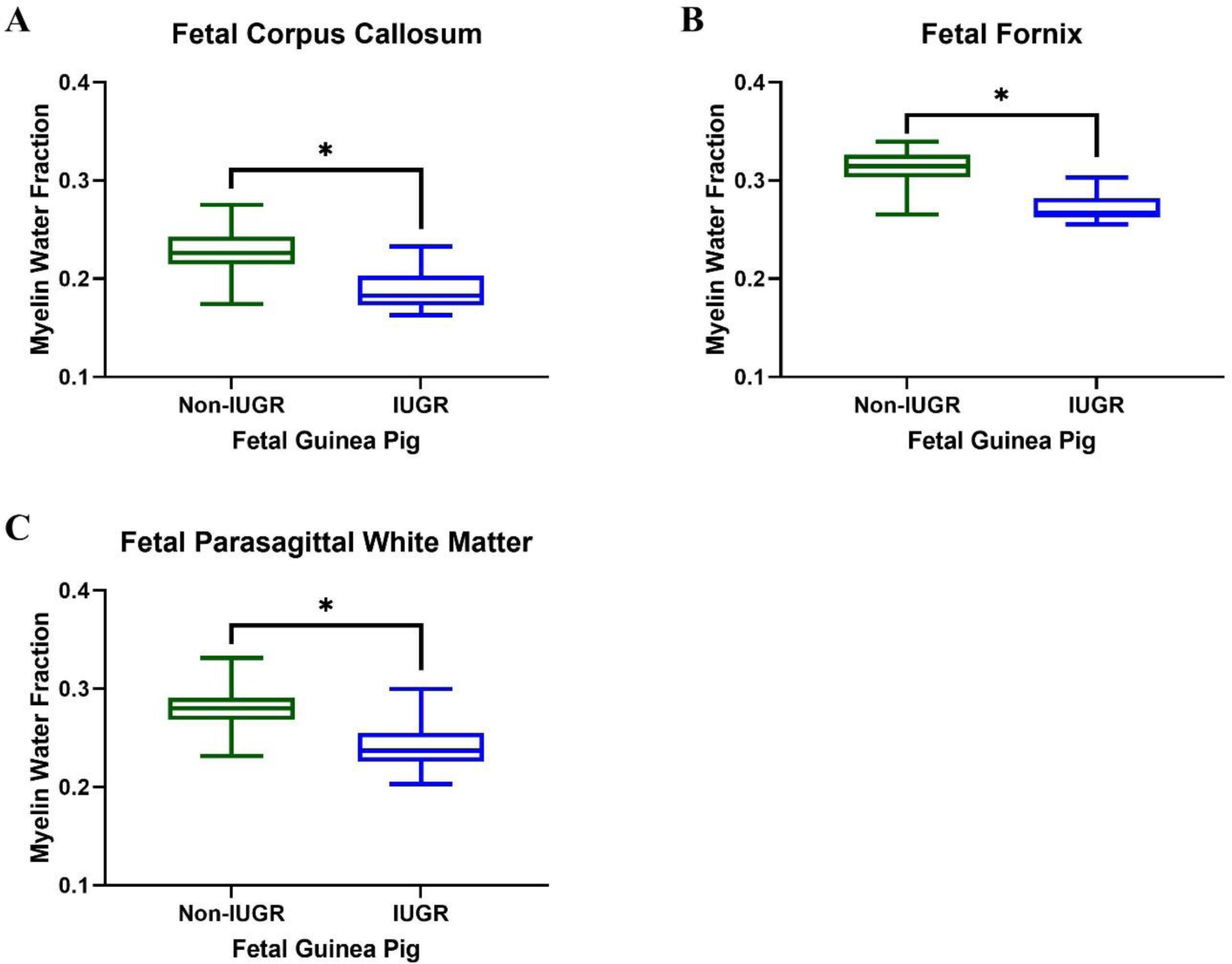
Box-and-whisker plot comparing the mean MWF of non-IUGR and IUGR fetal guinea pigs in the **(A)** corpus callosum, **(B)** fornix, and **(C)** parasagittal white matter. The guinea pigs comprising the IUGR group met three or more of the five IUGR criteria. The non-IUGR group’s guinea pigs met two, one, or zero of the five IUGR criteria. In all three regions, the mean MWF of the IUGR group was significantly lower than the mean MWF of the non-IUGR group, indicated by the black asterisk (*) [*p* < 0.05].

**Figure 3.**
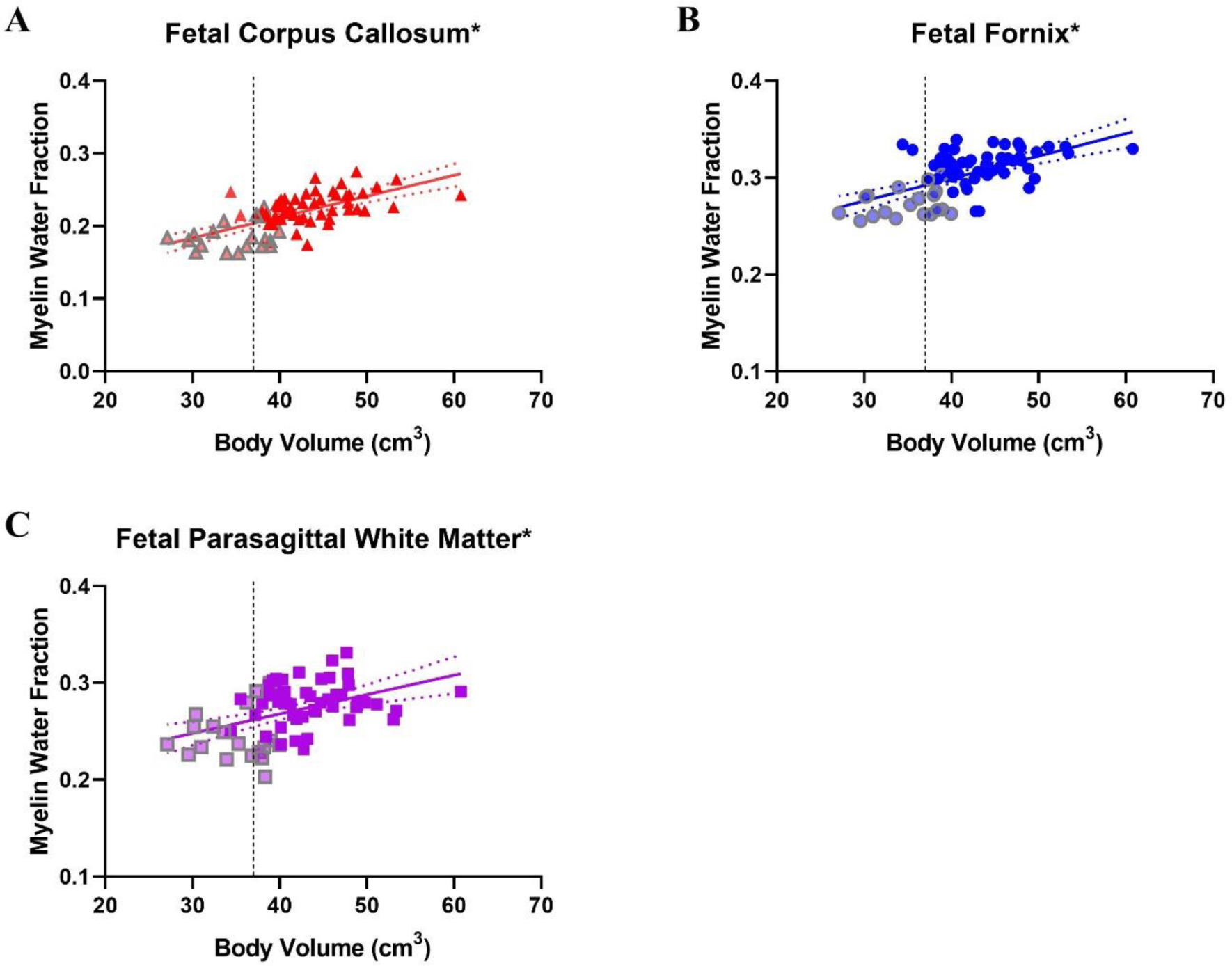
Linear regression line and 95% confidence intervals between body volume (BV) and myelin water fraction (MWF) of fetal guinea pigs’ **(A)** corpus callosum (CC) [red triangles], **(B)** fornix (FOR) [blue circles], and **(C)** parasagittal white matter (PSW) [purple squares]. The black dotted vertical line in all three graphs indicates the cut-off point for IUGR classification of a BV ≤ 37 cm^3^. The red triangles, blue circles, and purple squares that are semi-transparent with a grey outline represent the MWF of the CC, FOR, and PSW, respectively, of the fetal guinea pigs with IUGR as determined by the IUGR criterion. The coefficient of determination (R^2^) of BV with MWF of the three fetal regions are as follows: 0.42 (CC), 0.35 (FOR), and 0.20 (PSW). A black asterisk (*) indicates that the slope of the line is significantly different from zero as determined by an F-test (*p* < 0.05).

**Figure 4.**
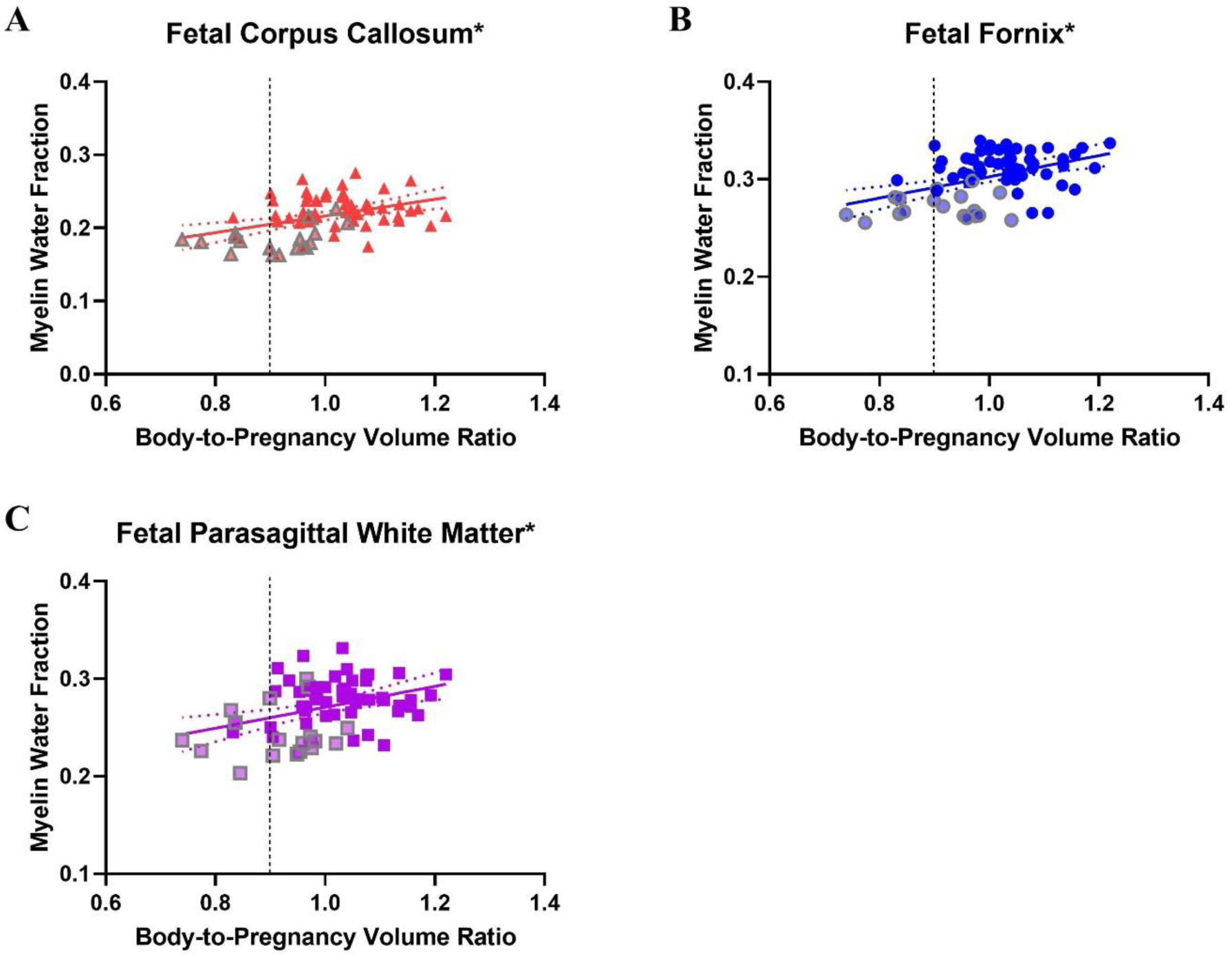
Linear regression line and 95% confidence intervals between body-to-pregnancy volume ratio (BPrVR) and myelin water fraction (MWF) of fetal guinea pigs’ **(A)** corpus callosum (CC) [red triangles], **(B)** fornix (FOR) [blue circles], and **(C)** parasagittal white matter (PSW) [purple squares]. The black dotted vertical line in all three graphs indicates the cut-off point for IUGR classification of a BPrVR ≤ 0.9. The red triangles, blue circles, and purple squares that are semi-transparent with a grey outline represent the MWF of the CC, FOR, and PSW, respectively, of the fetal guinea pigs with IUGR as determined by the IUGR criterion. The coefficient of determination (R^2^) of BPrVR with MWF of the three fetal regions are as follows: 0.17 (CC), 0.20 (FOR), and 0.14 (PSW). A black asterisk (*) indicates that the slope of the line is significantly different from zero as determined by an F-test (*p* < 0.05).

**Figure 5.**
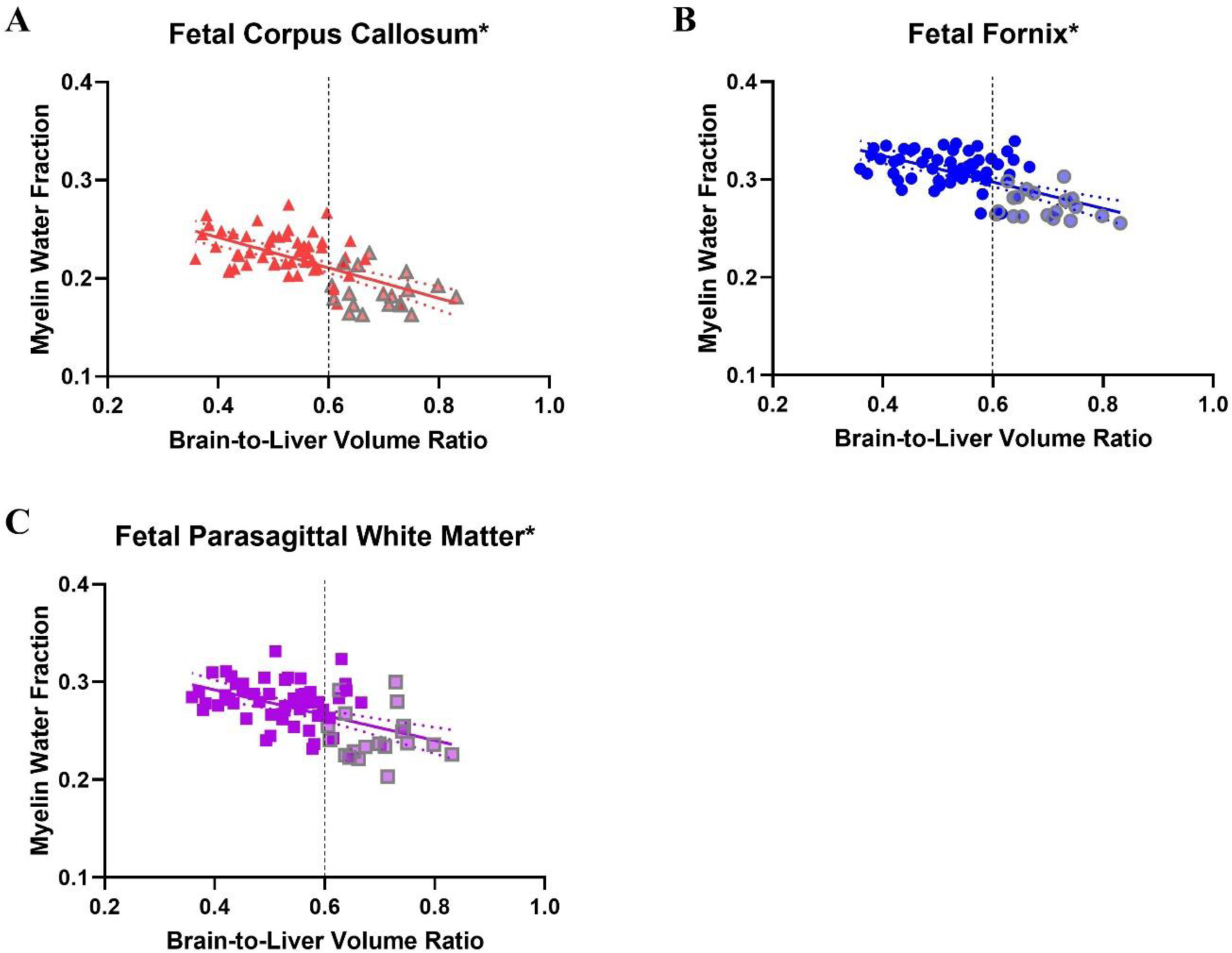
Linear regression line and 95% confidence intervals between brain-to-liver volume ratio (BLVR) and myelin water fraction (MWF) of fetal guinea pigs’ **(A)** corpus callosum (CC) [red triangles], **(B)** fornix (FOR) [blue circles], and **(C)** parasagittal white matter (PSW) [purple squares]. The black dotted vertical line in all three panels indicates the cut-off point for IUGR of a BLVR ≥ 0.6. The red triangles, blue circles, and purple squares that are semi-transparent with a grey outline represent the MWF of the CC, FOR, and PSW, respectively, of the fetal guinea pigs with IUGR as determined by the IUGR criterion. The coefficient of determination (R^2^) of BLVR with MWF of the three fetal regions are as follows: 0.40 (CC), 0.38 (FOR), and 0.27 (PSW). A black asterisk (*) indicates that the slope of the line is significantly different from zero as determined by an F-test (*p* < 0.05).

**Figure 6.**
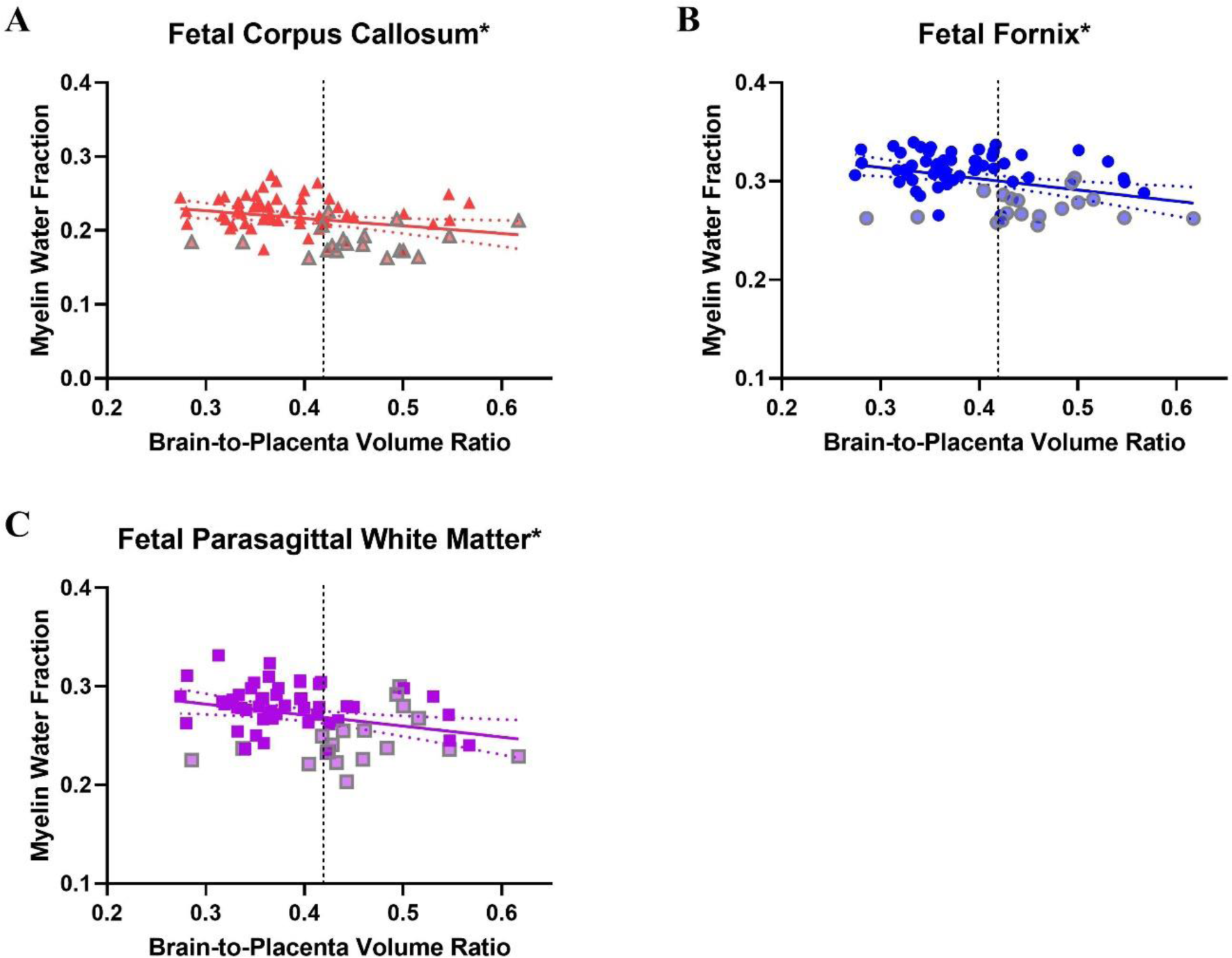
Linear regression line and 95% confidence intervals between brain-to-placenta volume ratio (BPlVR) and myelin water fraction (MWF) of fetal guinea pigs’ **(A)** corpus callosum (CC) [red triangles], **(B)** fornix (FOR) [blue circles], and **(C)** parasagittal white matter (PSW) [purple squares]. The black dotted vertical line in all three panels indicates the cut-off point for IUGR) classification of a BPlVR ≥ 0.42. The red triangles, blue circles, and purple squares that are semi-transparent with a grey outline represent the MWF of the CC, FOR, and PSW, respectively, of the fetal guinea pigs with IUGR as determined by the IUGR criterion. The coefficient of determination (R^2^) of BPlVR with MWF of the three fetal regions are as follows: 0.08 (CC), 0.13 (FOR), and 0.09 (PSW). A black asterisk (*) indicates that the slope of the line is significantly different from zero as determined by an F-test (*p* < 0.05).

**Figure 7.**
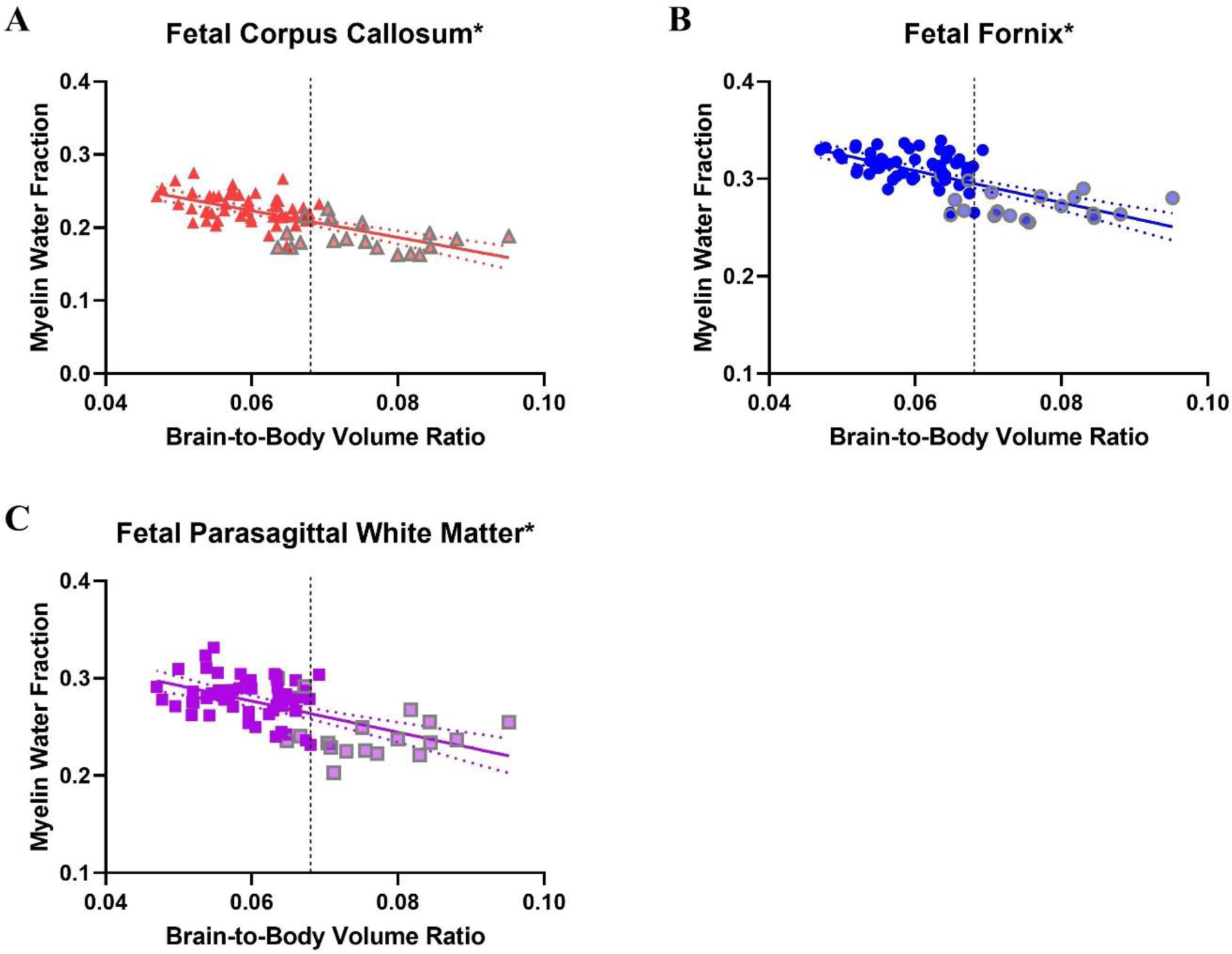
Linear regression line and 95% confidence intervals between brain-to-body volume ratio (BBVR) and myelin water fraction (MWF) of fetal guinea pigs’ **(A)** corpus callosum (CC) [red triangles], **(B)** fornix (FOR) [blue circles], and **(C)** parasagittal white matter (PSW) [purple squares]. The black dotted vertical line in all three panels indicates the cut-off point for IUGR classification of a BBVR ≥ 0.068. The red triangles, blue circles, and purple squares that are semi-transparent with a grey outline represent the MWF of the CC, FOR, and PSW, respectively, of the fetal guinea pigs with IUGR as determined by the IUGR criterion. The coefficient of determination (R^2^) of MWF of the three fetal regions with BBVR are as follows: 0.45 (CC), 0.47 (FOR), and 0.34 (PSW). A black asterisk (*) indicates that the slope of the line is significantly different from zero as determined by an F-test (*p* < 0.05).

**Table 3.** Linear Regression Comparison Between Weight & Volume Markers for Intrauterine Growth Restriction Determination.

| Marker Comparison | Slope ± SEM | Y-Intercept ± SEM | *P* value of slope | R<sup>2</sup> |
| --- | --- | --- | --- | --- |
| Body Weight (g) v. Body Volume (cm<sup>3</sup>) | 0.41 ± 0.03 | 2.32 ± 2.67 | < 0.05 | 0.96 |
| Body-to-Pregnancy Weight Ratio v. Volume Ratio | 0.97 ± 0.07 | 0.03 ± 0.07 | < 0.05 | 0.99 |
| Brain-to-Liver Weight Ratio v. Volume Ratio | 0.97 ± 0.01 | 0.02 ± 0.0 | < 0.05 | 0.99 |
| Brain-to-Placenta Weight Ratio v. Volume Ratio | 0.49 ± 0.06 | 0.13 ± 0.03 | < 0.05 | 0.5 |
| Brain-to-Body Weight Ratio v. Volume Ratio | 2.4 ± 0.16 | 0.0 ± 0.0 | < 0.05 | 0.8 |

Note. – SEM = standard error of the mean.

## DISCUSSION

This study quantified and compared MWF in the brains of fetal guinea pigs with and without IUGR. In all three regions analyzed – CC, FOR, and PSW – the MWF was significantly lower in the guinea pigs with IUGR. This study also determined the correlation of the IUGR markers of BV, BPrVR, BLVR, BPlVR, and BBVR with the MWF of the three analyzed regions. In all regions, BBVR was found to have the strongest correlations with MWF, while BPlVR had the weakest correlations with MWF.

Among the non-IUGR and IUGR fetuses, the FOR had the highest MWF, with the PSW being second and the CC having the least. These results are consistent with previously published results where MBP staining was the highest in the FOR, second in the PSW, and the lowest in the CC in guinea pigs of the same gestational age, supporting our results (29). In guinea pigs, myelination begins in the first half of gestation in the FOR and the second half in the CC (30). The largest difference in mean MWF between non-IUGR and IUGR guinea pigs was in the CC, followed by the PSW and then the FOR.

Like the feasibility study, this study employed mcDESPOT as the technique for MWI. Hence, this study’s imaging protocol was based on the protocol in the feasibility study, with the main difference being a decrease in voxel size to 0.6 mm^3^ from 0.7 mm^2^ (11). Since this study looked at only fetal brains as opposed to both fetal and maternal brains, the decrease in the field of view to only the uterus allowed increased resolution and improved identification of fetal brain structures. Consequently, we could identify and include the analysis for the PSW, whose myelin content is affected and may have been altered in IUGR as validated by histology (29), as opposed to only analyzing the CC and FOR, which was the case in the feasibility study (11). Since MWI was already conducted in fetal guinea pigs, this study continued to use this animal model, especially since the decrease in brain myelin content from IUGR was previously demonstrated in fetal guinea pigs of the same gestational age (11,29).

In previous animal studies, IUGR has been induced using models of maternal malnutrition, chronic hypoxia, and endocrine alterations to mimic the human conditions associated with an IUGR outcome (31). Another model successfully used in other studies is spontaneous IUGR, where IUGR occurs naturally and without artificial intervention. Guinea pigs can demonstrate spontaneous IUGR due to their variable and large litter sizes; this model produces fetuses with asymmetric IUGR that experience neonatal catch-up growth, which is seen in IUGR humans (15,32). Hence, this study opted for the spontaneously occurring IUGR model, with cases of placental insufficiency occurring naturally in the population similar to humans.

BW, BPrWR, BLWR, BPlWR, and BBWR have been previously used in animal studies as markers for IUGR (2,24–27), though generally in isolation and no studies have reported using multiple measures of IUGR. Instead of using weight markers, the current study employed the volume equivalent of these markers for clinical translation purposes. Since IUGR fetuses are classified as weighing below the 10^th^ percentile, BV was used as a marker for IUGR. BPrVR is also used because, in most pregnancies with IUGR fetuses in our sample, not every fetus within the pregnancy is growth-restricted. Among the remaining three volume markers, the common denominator is the brain. IUGR fetuses have a higher value for these markers than non-IUGR fetuses due to a phenomenon called brain sparing. In situations of hypoxia and nutrient deprivation, such as in IUGR, the fetus redistributes its cardiac output to maximize oxygen and nutrient supply to the brain (33). In the IUGR situation, evidence suggests this occurs as peripheral vascular beds vasoconstrict and cerebral arteries vasodilate so that the cardiac output favours the left ventricle. Since the left ventricle supplies blood to the upper body and brain, blood supply towards the brain is enhanced to preserve brain growth while supply to the organs in the lower half, such as the liver, is reduced (34,35). Consequently, the brain is proportionally larger than other organs in IUGR fetuses than in non-IUGR fetuses. As for the placenta, it is likely to be proportionally smaller in volume in IUGR since most cases of IUGR are due to placental insufficiency (5).

As previously mentioned, the fetal MWF of guinea pigs with IUGR was significantly lower than control guinea pigs in all three regions. Similar results have been observed in humans and guinea pigs, as validated by histology (7,29). The mechanism behind the IUGR-related myelin reduction is likely due to the often reported chronic hypoxia and oxidative stress associated with IUGR (36). Both effects are known to inhibit oligodendrocyte differentiation and, hence, myelination. Oligodendrocyte differentiation is controlled by a combination of inductive and repressive factors, with the main one being bone morphogenetic proteins (BMPs) (37). Of the 20 BMPs, BMP4 has the most significant role in inhibiting oligodendrocyte differentiation, especially as oligodendrocyte precursors are particularly vulnerable to oxidative stress due to their low endogenous glutathione levels, high iron content, and high oxidative metabolism (37). Elevated BMP4 signalling has been demonstrated in animal models of IUGR (38). Thus, it is likely that increased BMP4 signalling due to the hypoxic environment of IUGR leads to a decrease in oligodendrocyte differentiation and, hence, myelination.

Although the size markers for IUGR are used clinically, they are not always perfect in identifying all true growth-restricted fetuses. IUGR often manifests in small for gestational age fetuses that weigh below the 10^th^ percentile; however, some fetuses are genetically programmed to be small and, hence, are just small for gestational age but not growth-restricted. Conversely, some fetuses above the 10^th^ percentile for weight are not growing to their prescribed genetic potential and are growth-restricted. Although not meeting the classical definition of IUGR, this group is still negatively impacted neurologically, as evidenced by Burger *et al.,* where birth weight positively correlated with educational performance at 12 years of age (39). The ratios involving the brain are better identifiers for IUGR than just size alone due to brain sparing, but they are also not perfect markers for IUGR. One of the benefits of using MWF as a marker for IUGR is that it is a functional marker. As previously mentioned, one of the neurological consequences of IUGR is a reduction in brain myelin content. Since myelin content is independent of fetal size, myelin content should be reduced in IUGR fetuses whether the fetal weight falls below the 10^th^ percentile. The use of MRI, in conjunction with ultrasound, to assess fetal development is increasing in frequency (40), allowing the IUGR-related reduction in myelin content to be quantified along with the already established size markers. In short, the established size markers for IUGR are accessible and work well to identify most growth-restricted fetuses. However, we can use functional markers like MWF alongside size markers to identify all true IUGR fetuses.

### Limitations

The study had multiple limitations, with the first being partial volume affecting our MWF measurements. Due to the small features of the analyzed regions, a possibility exists that the ROI placement extended outside the desired region due to its small size. Although the number of ROIs placed in each region varied between animals to ensure the ROIs were only being placed in the desired region, the limitation was likely not completely avoided. Furthermore, this study lacked inter-rater reliability as only one individual conducted the ROI placement and the tissue segmentations. Another limitation is the potential motion between the T_2_-weighted acquisition used for ROI placement and the DESPOT_1/2_ acquisitions used to generate the MWF maps. Hence, the ROI may not have occupied the same voxels between the T_2_-weighted image and the MWF map. The study only looked at the development of myelin in IUGR late in gestation; the next step would be to image earlier in gestation and determine whether IUGR-related reductions can be identified and assessed earlier in gestation. Although the acquisition time for the DESPOT_1/2_ volumes was reduced compared to the feasibility study (11), the acquisition times are still substantially long. They must be dramatically reduced when transitioning to human fetal imaging, especially since using anaesthesia for human imaging would not be an option.

## Conclusion

This study showed that MWF is reduced in the CC, FOR, and PSW in fetal guinea pigs with IUGR compared to non-IUGR fetuses. The correlations between each IUGR marker and MWF were significant in all three regions, as BBVR and BPlVR had the strongest and weakest correlations with MWF, respectively.

## Acknowledgements

The authors would like to thank Paige Allen for their help with the linear mixed model analysis of the data. The authors would also like to thank Brian Sutherland for their help with animal tissue collection.

## Notes

**Grant Support:** NSERC Discovery Grant (RGPIN-2019-05708) NSERC Post-Graduate Scholarship – Doctoral

### Competing Interest Statement

The authors have declared no competing interest.

